# Deep Learning-Enhanced TopoStats for the Automated Quantification of DNA and Complex Biomolecular Structures

**DOI:** 10.64898/2026.05.06.723223

**Authors:** Sylvia Whittle, Tobias A. Firth, Max Gamill, Laura Wiggins, Neil Shephard, Toby Allwood, Thomas E. Catley, Alice L. B. Pyne

## Abstract

Atomic force microscopy (AFM) enables nanometre-scale, label-free imaging of biomolecules and surfaces under near-native conditions, yet quantitative analysis of AFM data remains limited compared to other bioimaging modalities. This limitation largely arises from the absence of open, automated tools capable of addressing AFM-specific artefacts, data formats, and topographical outputs. Here, we present the latest version of TopoStats, an open-source Python package for automated and quantitative AFM image analysis, developed as a deep-learning enabled advancement of our original TopoStats software to support more complex samples and richer molecular characterisation. The pipeline integrates all key processing stages, including image flattening and noise correction, object detection and segmentation, morphometric feature extraction, and strand tracing with topological classification. Designed for accessibility and reproducibility, TopoStats adheres to the FAIR for Research Software (FAIR4RS) principles and provides configurable workflows adaptable to diverse biological samples. Combining high-resolution AFM and our analysis pipeline allows the quantification of subtle structural changes within a heterogeneous sample set, revealing properties not accessible with other structural biology techniques. We demonstrate the effectiveness of our pipeline to differentiate between plasmids with both different topology and sequence, by extracting meaningful quantitative descriptors that distinguish the samples with statistical significance. Collectively, these developments establish TopoStats as a versatile framework for high-throughput, quantitative AFM analysis, advancing AFM from a fundamentally qualitative visualisation technique toward a quantitative analytical tool.

## 1. Introduction

Advances in AFM technology have made routine high-resolution imaging of biomolecules and polymers accessible to the wider biophysics community. Commercial systems with a stable, high signal-to-noise ratio provide users with the ability to image with sub-nanometre lateral and picometre axial resolution, allowing for label-free imaging in near-native conditions. The introduction of ‘tapping mode’ allowed for the first image of DNA to be resolved by AFM^1,2^, with later technical developments allowing resolution of the major/minor grooves to be readily attainable^3^. Similarly, AFM has been used to characterise the structure of pathological protein fibrils associated with Alzheimer’s disease and the heterogeneity of actin filament substructure demonstrating AFM as a viable platform for diverse biological research applications^4,5^. Despite these strengths, wide-scale adoption of the technology faces a distinct challenge; the availability of robust and accessible quantitative analysis tools that remove reliance on coarse shape metrics and/or qualitative interpretation.

The development of these tools is hindered by the unique nature of AFM data. The community is frequently challenged by proprietary file formats instituted by the manufacturers, which limit the use of external analysis workflows. Furthermore, AFM data presents specific processing hurdles that cannot be solved using tools designed for non-surface techniques. AFM requires specialised preprocessing to correct for specific artefacts such as row misalignment, scanner drift and tip convolution, and feature extraction must account for surface topography rather than intensity-based contrast common in optical methods. Whilst a host of powerful AFM analysis tools exist e.g., Gwyddion^6^, NanoLocz^7^ and WSxM^8^, there isn’t a clear standardised workflow for automated, high-throughput polymer analysis that can generate detailed statistics on molecular conformation.

This is in contrast to the broader bioimaging community who have seen a shift towards automated workflows for batched quantitative analysis and open-source software. Bioimaging encompasses a diverse array of microscopy approaches, each producing vastly different data types. While manufacturers often provide proprietary analysis software, the community repeatedly returns to open-source, accessible software such as Fiji^9^, CellProfiler^10^, RELION^11^ and Napari^12^, all of which provide flexible, modular environments for basic pre-processing to the most complex analyses. The open-source and modular nature of these tools also enable researchers to adapt and build upon existing workflows for specialised samples and to develop new analysis methods. Machine-learning based tools like DeepImageJ^13^ have added a new dimension to image analysis by providing convenient integration of advanced segmentation and classification into workflows. The persistent use of open-source, community-driven workflows instead of bespoke tools by the bio-imaging community, reflect its motivation for reproducible and accessible software.

In response to the need for a unified, accessible, and quantitative tool for AFM image analysis, we present TopoStats v2.0.0, an enhanced successor to TopoStats^14^ that supports more complex sample types and enables richer biomolecular characterisation. TopoStats v1.0.0 provided 20 metrics describing object or molecule geometry, whereas TopoStats v2.0.0 provides a total of 68 metrics across several additional categories. TopoStats now provides an automated pipeline comprising five key stages **(Figure 1)**: 1) image loading via AFMReader, 2) preprocessing, 3) deep-learning augmented object detection, 4) morphological analysis for objects of interest, 5) molecular tracing and analysis and was developed in accordance with the FAIR for Research Software (FAIR4RS) principles^15^.

**Figure 1.**
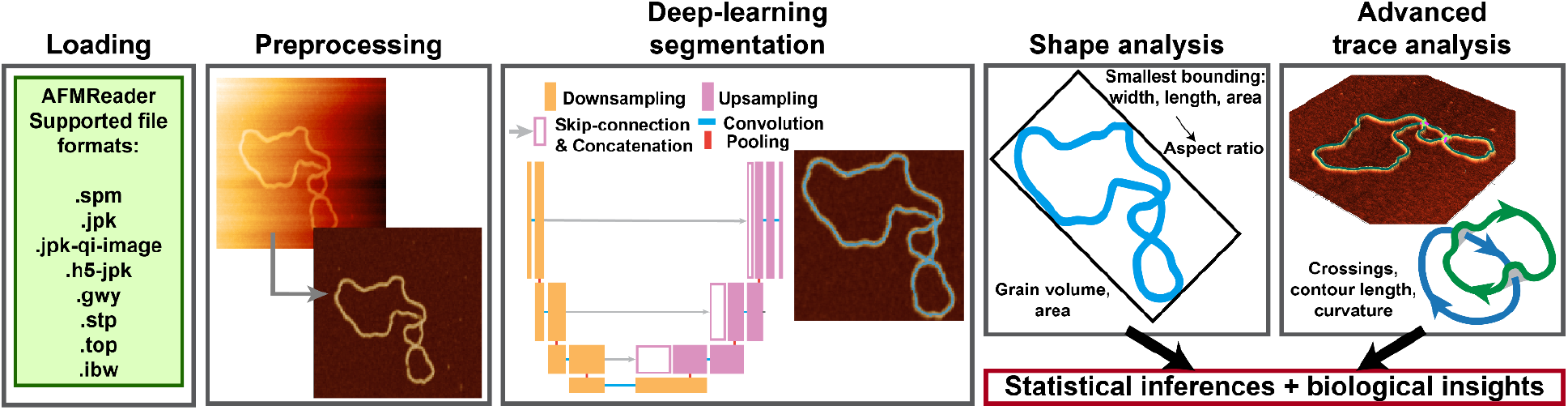
TopoStats processing and quantification pipeline. TopoStats takes raw AFM data, preprocesses and segments objects of interest through a combination of thresholding and deep learning segmentation. An advanced DNA trace analysis pipeline then quantifies the complex conformations of molecules, which are able to be used for statistical analysis.

To highlight the full capabilities of this workflow we use DNA as an exemplar biomolecular system with tuneable length and topological complexity. We show that we can segment individual molecules and trace their paths, providing quantitative measurements of their length, flexibility and degree of self-interaction, driven by sequence or topological changes. DNA topology is fundamental to numerous biological processes and interactions, for example, supercoiling and DNA secondary structures are known to regulate transcriptional activity^16–20^, and protein binding events^21–23^. The conformational variability of DNA often makes measurements of these effects difficult as they cannot be described as a single structural class. Single molecule AFM measurements have been shown to be an effective way to image and describe the structure of DNA, with polymeric analysis showing indicators of metrics such as contour and persistence length^24–26^. However, these measurements often require manual intervention or are based on in-house developed code and are therefore difficult to compare between laboratories and samples. Here we demonstrate TopoStats as a powerful tool for the quantification of biomolecules between datasets and across a range of samples, showing how the number of crossings and curvature vary for DNA in different topological states. We then explore how these metrics are altered when the DNA contains an AT-rich tract, compared to a telomeric repeat insert. Overall, TopoStats provides a robust, open-source pipeline for the quantification of AFM data across a wide range of biomolecular samples.

## 2. Results

### 2.1. Standardised AFM file loading with the AFMReader Python package

Loading and extracting data from AFM image files is notoriously difficult due to competing manufacturer file types, standards and insufficient documentation. While existing analysis software^6,8^ can handle these files, they are limited in their ability to interface with modern Python libraries which restricts downstream analysis. There is some community effort to provide tools to make these formats more accessible, with libraries like pySPM^27^ and igor2^28^, however they are not unified and their outputs have to be handled on a case-by-case basis. We have developed AFMReader^29^, a Python package that unifies support for multiple AFM file formats within a single framework, using both existing, community made libraries and bespoke loading apparatus. This is designed with interoperable open-source code that is thoroughly tested, incorporating comprehensive documentation and community involvement. AFMReader currently supports the most widely used file types from major AFM manufacturers and, using the HDF5 file structure, provides a unified output to enable the development of processing scripts and analysis software that are agnostic to instrument or file type (**Supplementary Table 1**). It functions both as an independent Python package and as an integrated module within TopoStats, extending the software’s applicability across all supported AFM file formats.

### 2.2 Automated pre-processing of AFM images to enable quantitative analysis

Once loaded, raw AFM images need to be pre-processed prior to downstream analysis to remove AFM-specific imaging artefacts^30^. TopoStats facilitates a series of configurable processing steps (**Supplementary Figure 1**) including tilt correction for correcting image slope, row alignment to remove scan-line artefacts, and an algorithm to remove scars - long, thin horizontal streaks in images due to the AFM tip erroneously skipping across the surface. Finally, the image data can be shifted so the background has a height of 0, allowing for reliable comparison of molecular heights between objects in multiple images.

During slope and row correction, issues can occur when surface features interfere with background estimation. In such cases, the algorithm may incorrectly interpret object regions as part of the background, causing the slope removal to perform poorly and leading to rows containing objects being over-corrected. This results in those rows appearing darker in the processed image and the object heights being underestimated **(Supplementary Figure 2A)**. To mitigate this effect, two improvements were introduced. First, the quantile used to estimate the background level for each row can now be user-configured, allowing finer control over how the algorithm defines the background height. Second, the preprocessing pipeline has been divided into two sequential stages. The initial stage performs standard image flattening and correction procedures to produce a preliminary image. The secondary stage then identifies non-background regions using a user-defined automatic thresholding method, masks these regions, and performs a second round of flattening restricted to background pixels. This two-step approach yields a more reliable and uniform image flattening, even in datasets containing a high density of surface features **(Supplementary Figure 2B)**. As well as improving the batch processing capability of TopoStats, this advance in flattening enables a higher sample throughput by use of higher surface concentrations of biomolecules, a factor that is typically limiting in single-molecule approaches.

### 2.3. Automated segmentation and feature extraction from AFM images

#### 2.3.1. Classical segmentation methods

To extract information on molecular conformation, objects of interest must first be identified through segmentation, to isolate the regions intended for measurement. In AFM topographs, objects of interest typically differ in height from the background, making height-thresholding a reasonable and effective approach for segmentation (**Supplementary Figure 3A**). We implemented three commonly used methods for height thresholding: Otsu, standard deviation-based, and absolute thresholding. The Otsu thresholding method estimates the optimal cutoff value for a bimodal height distribution, generating a mask of all pixels above this threshold. This approach is particularly effective when objects of interest exhibit similar, well-defined heights relative to the background. The absolute thresholding method allows direct specification of the threshold in nanometres, making it useful when the approximate height of the target structures is known or a precise cutoff is desired. The standard deviation method defines the threshold as the mean height of the grains plus a user-configurable multiple of the standard deviation, providing flexibility for cases where object heights are unknown or vary widely according to a normal distribution.

#### 2.3.2. Segmentation clean-up steps

After an initial segmentation mask is generated (**Supplementary Figure 3B**), cleanup is required to ensure that only objects of interest are retained in the segmentation mask for analysis. Objects that intersect the edge of the image cannot be reliably measured since an indeterminate amount of the object lies outside the image bounds, and so these objects are optionally removed (**Supplementary Figure 3C, blue**). Furthermore, objects of interest may clump together, or there may be contamination in the image that could interfere with analysis. These can be removed using configurable area thresholds, removing objects that are smaller or larger than the expected object size, (**Supplementary Figure 3D, purple**). The result is a mask that ideally, depicts just the objects intended for downstream analysis and quantification.

#### 2.3.3. Deep-learning methods for improved segmentation

The quality of downstream analysis is directly tied to the accuracy of sample segmentation, making the generation of an accurate mask essential. Conventional height-thresholds, while effective for simple, well-defined samples, often struggle when applied to heterogeneous data where objects of interest vary in height, overlap, or display only subtle topographical contrast with the background. The increasing use of deep learning models^31,32^ has driven huge advances in segmentation tools^33–35^ that, in some cases, exceed the capabilities of traditional approaches. Having demonstrated the power of deep learning models to accurately segment complex biological species e.g., catenanes whilst preserving the crucial structural details often obscured during traditional height thresholding (Holmes et al.^36^), we elected to integrate deep learning into the segmentation functionality of TopoStats. Using tools such as LabelStudio, users can label images, highlighting specific features of interest and use this to train a deep learning model. Once trained, users can add their bespoke TensorFlow model easily within TopoStats by linking their model to the standard configuration file where it will be used as part of the segmentation workflow. The implementation of deep learning models in TopoStats provides users with the tools to further refine their image analysis and enhance object segmentation for their unique sample^36^. An example of segmentation improvements from a Unet model trained on plasmid DNA is shown in **Figure 2**. Each region of interest is cropped, padded by a configurable amount, normalised and then run through the segmentation model to produce a mask that more faithfully preserves fine structural details of objects of interest that traditional methods often obscure.

**Figure 2.**
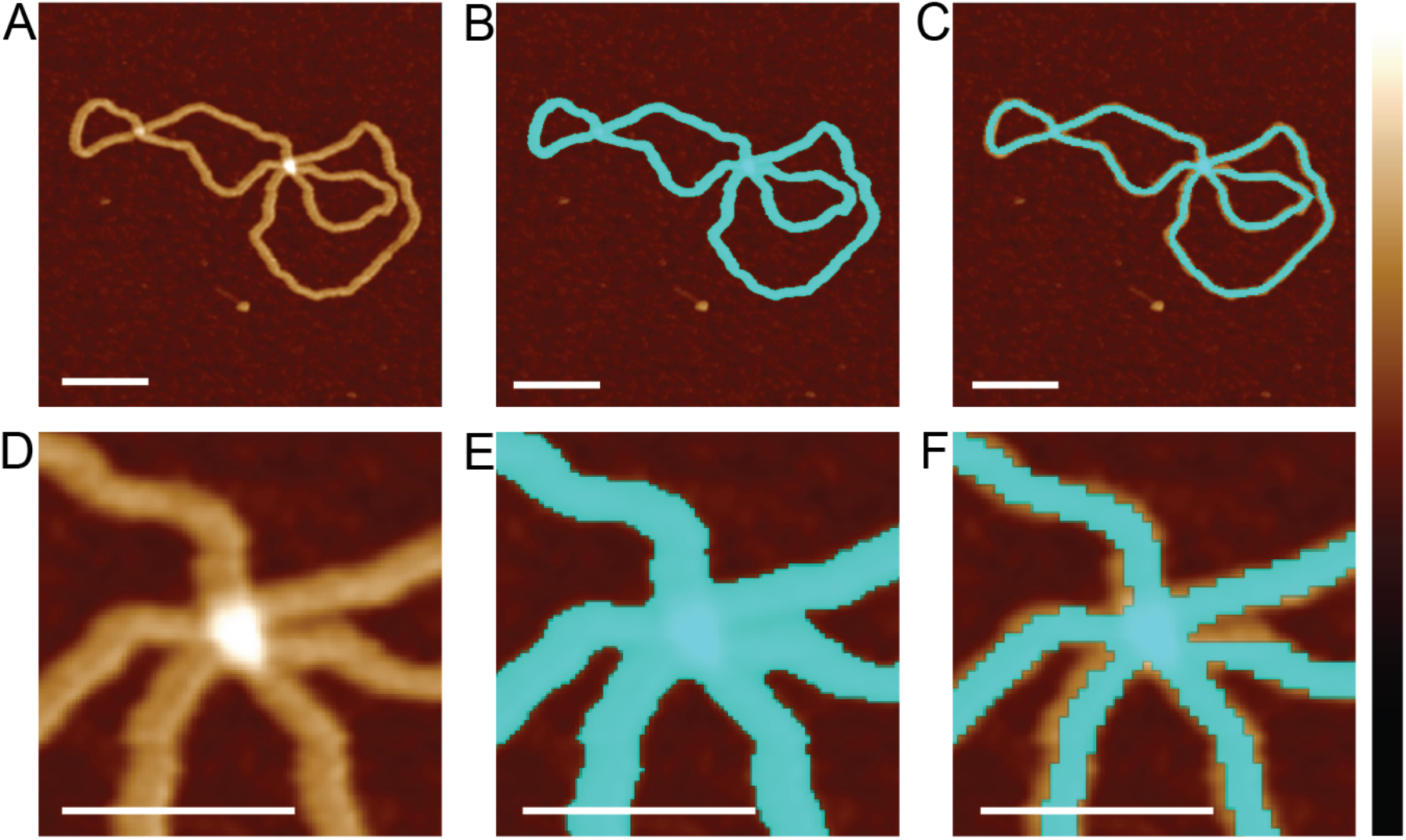
Deep learning to refine object segmentation. AFM image of a DNA molecule **(A)**. Traditional height-thresholded mask of the molecule **(B)**. U-Net segmentation of the molecule enables better masking **(C)**. Zoom-ins of a particularly difficult to segment crossing of the catenane highlighting the improvement due to the U-Net segmentation **(D-F)**. Scale bars: 50 nm. Height scale: −3 nm to +4 nm.

Once objects have been successfully segmented, their corresponding height-map pixels are used to calculate quantitative morphological statistics. We implemented a suite of scripts to generate a comprehensive set of statistics, including spatial and geometric information, such as the position, aspect ratio, feret diameters and bounding areas, and height-based information including height and volume (**Supplementary Figure 4**). These statistics are output into a human-readable CSV file indexed by image, folder name, and object number for easy identification and comparison between datasets.

### 2.4. An advanced skeletonisation pipeline for identification of molecular backbones

For filamentous samples such as DNA, fibrils or polymer chains, global morphological statistics can fail to fully describe conformational differences between objects. In these cases, it is often more informative to determine the molecular contour length or width and determine whether individual molecules adopt a linear or closed-loop conformation. While other strand tracing pipelines allow for calculation of these metrics^14,26^, they cannot accurately identify and trace self-interaction events, where two strands cross over each other. These events are important to resolve to understand how polymers are interlinked, and how often they cross one another. Correctly tracing strands as they pass over or under one another is essential for accurately describing molecular topology^36^ and enables the separation of overlapping or intersecting molecules. These metrics are a useful indicator of conformational state, for example when distinguishing relaxed and supercoiled DNA^36^, and can be used to quantify the effects of conformation-altering drugs^37^. We have developed a sophisticated tracing pipeline that harnesses the unique topographical information available from AFM images to produce reliable traces of molecular backbones, including at intersecting points.

#### 2.4.1. Optimising segmentation masks for skeletonisation

The first step is to generate a representative skeleton - a single-pixel-wide mask that accurately delineates the central axis of the segmentation mask (**Figure 3A,B**). As skeletonisation is applied directly to the mask, it is often necessary to pre-process masks, for example by smoothing edges or filling small holes, to minimise the formation or erroneous skeleton branches. To support this, we implemented two pre-processing options: 1) gaussian smoothing followed by re-binarising using Otsu thresholding and 2) binary dilation. Gaussian smoothing is effective for reducing noise in the mask whilst maintaining the shape of the grains. The second method, binary dilation, performs morphological dilation to smooth and connect adjacent regions of the mask which can help close small gaps and strengthen thin regions of a mask. Both methods yield a smoother mask edge and reduced instances of spurious branches in the skeletonised trace. In some instances, the trace can break as it passes through low regions of the molecule (for example a region of single-stranded DNA within a plasmid). To avoid unnecessary grain removal during data clean-up due to these incomplete traces, we implemented an end joining module that detects and joins nearby endpoints **(Supplementary Figure 5)**. This in turn allows gaps in segmentation masks to be retrospectively filled and the affected grain to re-enter the analysis workflow.

**Figure 3:**
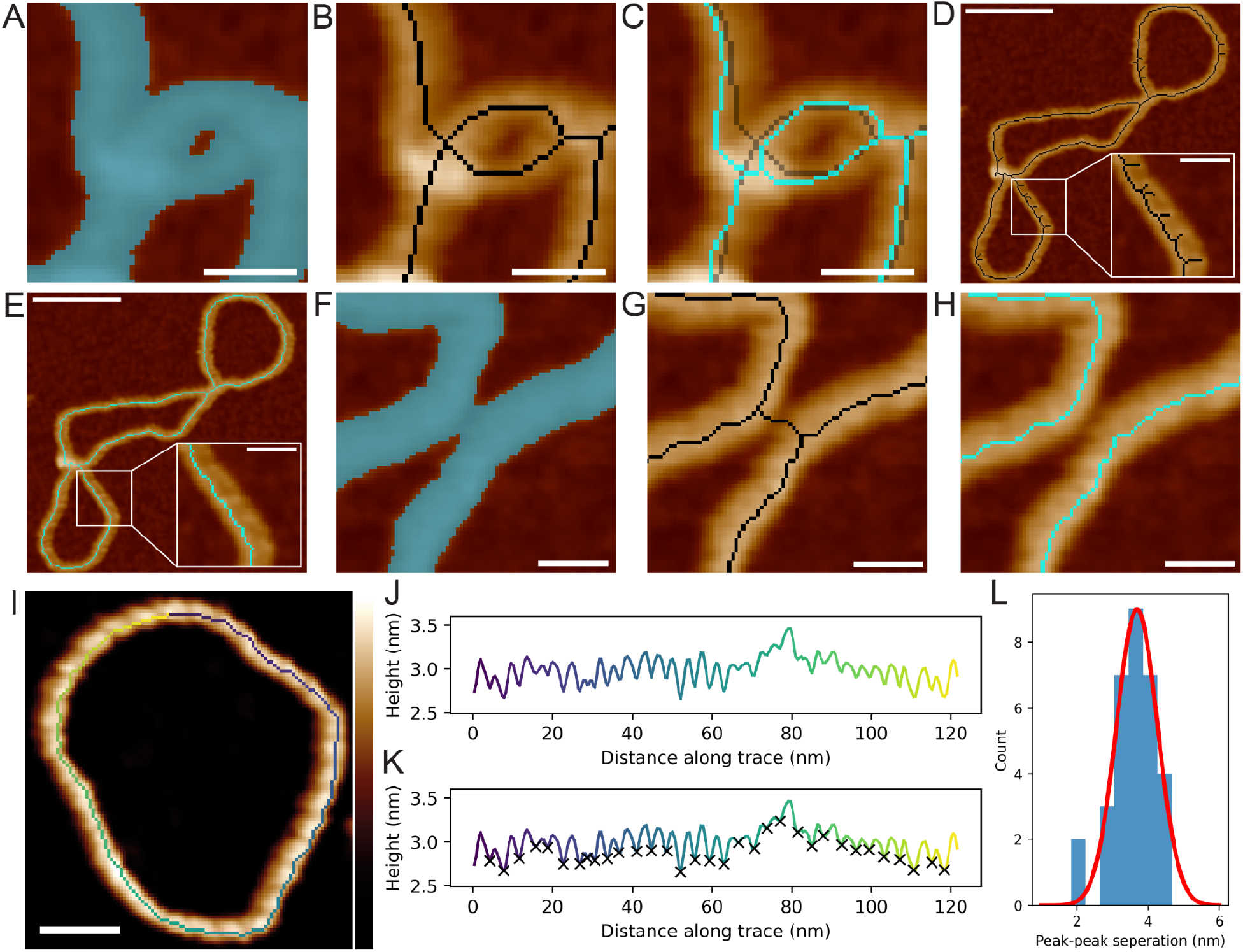
Advanced skeletonisation analysis removes errors allowing full plasmid traces to be obtained. Height biasing applied during the skeletonisation of a single pICOz molecule mask**(A)**pulls unrepresentative skeletons **(B)** towards higher and more representative regions of DNA strands **(C)**. Skeleton pruning of a single pICOz trace **(D)** removes erroneous trace branches **(E)**. Skeleton pruning based on height of branches between two close, but separate, pICOz strands removes the erroneous connecting branch **(G), (H)**. Accurate skeletonisation of minicircle DNA yields a height trace **(J)**, which allows the detection of the major grooves location **(K)**. Measurement of the major groove spacing, with a Gaussian fit in red **(L)**. Scale bars: A, B zoom-outs = 40 nm. A, B zoom-ins = 10 nm. C-H = 10 nm. I = 10 nm. Height scale: A-H = −3 nm to +4 nm, I = −0.5 to 4 nm.

#### 2.4.2. Height-biased skeletonisation optimises molecular traces

Following mask pre-processing, a skeleton representation of the object mask can be obtained using a traditional skeletonisation method such as the Zhang algorithm^38^. However, in AFM images, tip convolution introduces positional uncertainty meaning that the molecular backbone is best represented by the highest regions of polymeric strands. Traditional binary morphological skeletonisation methods do not account for the underlying height information produced by AFM, often resulting in skeletons that deviate from the true molecular backbone. We therefore developed a customised version of Zhang’s skeletonisation algorithm that biases the skeleton towards higher regions of strands. In each iteration, the algorithm identifies pixels eligible for deletion but removes only a user-defined fraction of them, preferentially retaining those corresponding to local height maxima. This approach guides the skeleton to remain centred on the highest pixels throughout the thinning process, producing a final skeleton that more closely follows the true molecular backbone thus providing a more reliable representation of the molecule (**Figure 3C**). Despite mask pre-processing, erroneous branches can still occur. To address this, we include a pruning algorithm that removes spurious branches based on their length and the underlying AFM height information (**Figure 3D,E**). This addition is motivated by the observation that erroneous branches often do not lie over regions of the molecule of interest but instead extend into background areas or bridge two regions across a background gap (**Figure 3F–H**). Branches essential for maintaining skeleton continuity are preserved to prevent fragmentation of a single molecule into separate traces. Full and accurate skeletonisation enables a host of advanced analysis features including determining the periodicity of surface features^39^. By taking the height trace along the backbone of the molecule (**Figure 3I,J**), it is possible to measure subtle height changes along a molecule with a high degree of accuracy; enabling us to measure the major groove spacing around a full DNA minicircle of 3.7 ± 0.6 nm **(Figure 3K,L**) which corresponds well to the known value of 3.5 nm for B-DNA^40^.

### 2.5. Crossing analysis for topological resolution of intersecting polymer strands

Beyond height tracing, accurate skeletonisation enables a pipeline which can determine the topology of DNA plasmids and more complex structures by identifying over- and underlying strands. Once a skeleton has been obtained and erroneous branches have been removed, crossing points can be identified as pixels in the skeleton that are connected to more than two neighbouring pixels, corresponding to locations where two strands intersect **(Figure 4A)**.

**Figure 4:**
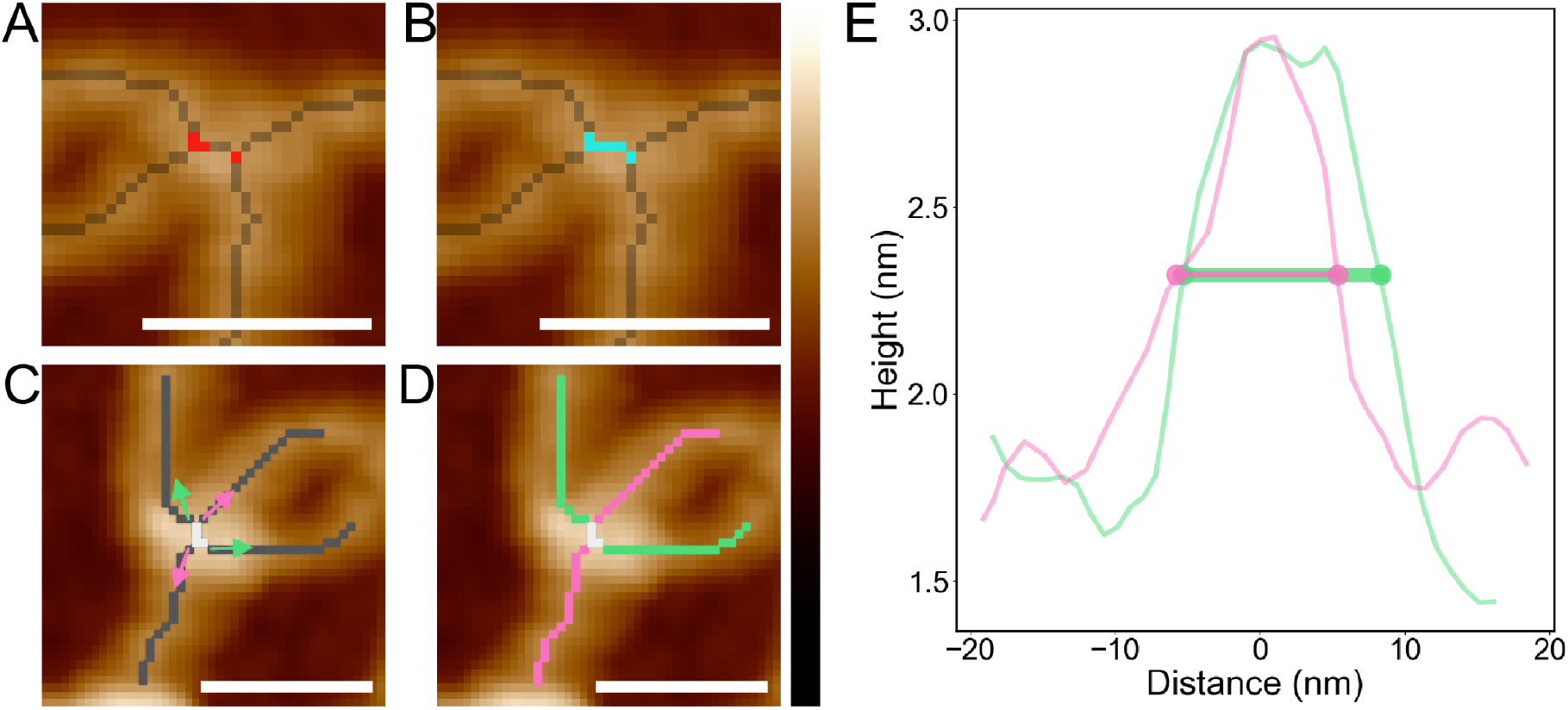
Detailed junction analysis enables tracing DNA through crossings. Merging of two separately identified junctions **(A)** into a combined junction **(B)**, enabling consideration of all four strands as part of one crossing. Vectors calculated from the strands **(C)** are used to connect similarly oriented strands to determine DNA paths through the crossings **(D). (E)** Consideration of the heights of each strand pair allows determining which strand pair lays over the other, by calculating which has a largest full-width-half-maximum height **(E)**. Scale bars: 40 nm. Height scale: −3 nm to +4 nm.

#### 2.5.1. Pairing branches at crossing points

Having identified these crossing points, the next step is to pair the emanating branches to achieve a continuous trace of the entire molecule. In some cases, what should represent a single intersection point is detected as multiple nearby crossing points. This effect arises from AFM tip convolution, which broadens the contact region between strands, extending the crossing area beyond its true dimensions (**Figure 4A**). Before determining how segments connect and which strand passes over or under, crossing points in close proximity are first merged and treated as a single node (**Figure 4B**). Once close nodes have been correctly consolidated, each crossing region is examined in turn, and the branches emanating from it are identified. Segment connectivity through each crossing is determined by analysing the angles formed by the branches emanating from the crossing centre. These angles are calculated by representing each branch as a single vector, obtained by following the branch for a user-defined distance and averaging the corresponding point coordinates (**Figure 4C**). Branches are then paired according to the closest matching angles, indicating the most likely continuation of the molecule through the crossing (**Figure 4D**). In some cases, crossings may contain an odd number of branches. In such instances, the algorithm either performs best-effort pairing, leaving one branch unpaired, or omits pairing entirely, based on user defined configuration parameters informed by sample morphology.

#### 2.5.2. Determining the stacking order of branches

The stacking order of overpassing and underpassing strands, i.e. which strand lies on top, and which lies below, is determined by analysing the height profiles of the strands as they pass through each crossing. Overlying and underlying DNA strands exhibit distinct height characteristics: the overlying strand gradually rises before reaching its peak height above the crossing, whereas the underlying strand remains relatively flat before showing a sharp increase as it passes beneath the other. This observation can be quantified by calculating the full width at half maximum (FWHM) of each strand’s height profile. The overpassing strand consistently exhibits a higher FWHM, while the underpassing strand has a lower value (**Figure 4E**). The difference between the FWHM values is used to assess the confidence in correctly identifying the over- and underpassing strands, with a larger difference indicating greater certainty in the classification.

### 2.6. Reconstruction of continuous DNA traces enables topological identification

Combining tracing and crossing analysis, TopoStats can reconstruct molecules as continuous traces. This is achieved by following connected pixels from a starting branch along the skeleton **(Figure 5A)** and through predetermined crossings to create a full, uninterrupted trace of the molecular contour (**Figure 5B,C**). In some cases, multiple molecules may be connected within the same grain, for example if two molecules are catenated (**Figure 5D-F**) or overlapping (**Figure 5G-I**), making it inaccurate to represent them as a single entity. To address this, the algorithm continues tracing unvisited branches and identifies them as separate molecules. This functionality is particularly valuable for tracing complex topological structures or to perform instance segmentation and separate analysis of distinct but overlapping molecules.

**Figure 5:**
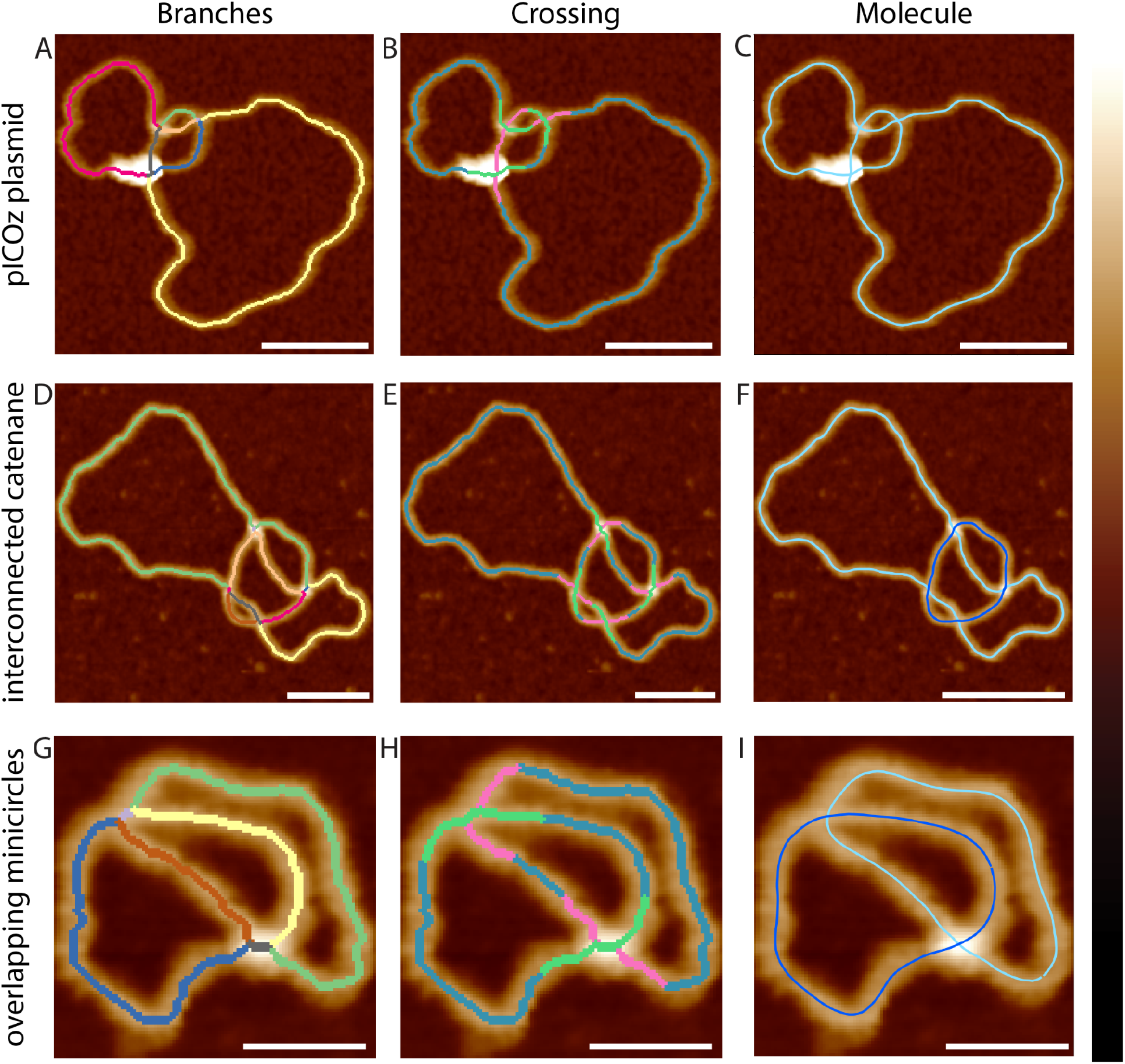
Trace vectorisation enables individual molecule identification. Splining and molecule identification from AFM images of a DNA plasmid exhibiting self-interactions **(A-C)**, an interconnected DNA catenane **(D-F)**, and two overlapping DNA minicircles **(G-I)**. Each of the skeleton branches obtained from the tracing are displayed in a different colour, **(A**,**D**,**G)**. Crossing determination is carried out with overlying strands marked in pink, and underlying strands in green, **(B**,**E**,**H)**. Smoothed trace of the plasmid using the rolling average method, **(C)**, the two interconnected molecules of the catenane have been automatically identified (small plasmid in dark blue and large plasmid in light blue), **(F)** and the two overlapping minicircles have been identified (each individual molecule spline is highlighted in dark or light blue), **(I)**. Scale bars: A-C 40 nm, D-F 80 nm, G-I 20 nm. Height scale: −3 nm to +4 nm.

#### 2.6.1. Trace smoothing to reconstruct the molecular backbone

Using pixel-based images for analysis inherently produces traces that are jagged, as each coordinate is constrained to the pixel grid, limiting smoothness to the resolution of the image. To overcome this, spline fitting can be applied, in which a mathematical curve is fitted through the trace points and resampled to generate a smoother trace. While effective in most cases, spline fitting requires careful parameter tuning to match the geometry of each molecule. If these parameters are not optimised then the spline can deviate significantly from the true molecular path, particularly around tight bends, yielding an unrepresentative trace where the trace ‘falls off’ the molecule and touches the background of the image. To address this limitation, we implemented an alternative smoothing method based on a rolling average which requires only a single, user-defined window size in nanometres (**Figure 5C,F,I**). This method is more reliable than splining, ensuring that traces do not stray from the molecule, and requires less optimisation by the user. Following smoothing, trace coordinates may be unevenly distributed with some regions dense in coordinates and others fairly sparse. To address this, we incorporated a resampling step that enforces uniform spacing of trace coordinates, with the rate of sampling able to be set by the user to increase or decrease the number of points as required for further analysis. The implementation of splining provides the foundation for measuring further tracing-based parameters that yield additional information to the grain-based parameters **(Supplementary Figure 4)**.

The implementation of splining provides the foundation for calculating additional tracing-based metrics that extend beyond shape descriptors such as aspect ratio. Using the generated spline we can calculate curvature; this parameter captures changes in molecule shape by measuring how sharply the trace bends locally. By quantifying the variation in curvature, we can determine changes in molecule conformation on the surface and make inferences about the underlying mechanics.

### 2.7. Accurate tracing metrics show significant differences between molecular topologies

#### 2.7.1. Supercoiled plasmids show a significant increase in number of self-crossings compared to nicked plasmids

Supercoiled DNA is known to have a more compact conformation due to superhelical stress which manifests as writhe, wherein two strands cross over themselves^40,41^. Conversely, nicked DNA has been shown to have a more open conformation, a consequence of a break in a single strand of the DNA backbone which releases the superhelical tension; nicked plasmids are less prone to self-crossing. We imaged both supercoiled and nicked pICOz (1277 bp) plasmids (n=52 supercoiled, n=34 nicked) and extracted grain- and trace-based statistics. When comparing the grain-based metric, aspect-ratio, we detected no differences between the supercoiled and nicked population **(Supplementary Figure 6A)**. For both supercoiled and nicked samples (**Figure 6A,B**), the mean contour length measured from the spline was close to the expected plasmid length of 434 nm, with a length of 417 ± 53 and 434 ± 83 nm for supercoiled and nicked respectively (**Figure 6C)**. As expected, crossing analysis revealed a significant change in the crossing distributions between the supercoiled and nicked plasmid populations, proving a potent metric for determining conformation and demonstrating the heterogeneity of the plasmids in each group (**Figure 6D,E**). The supercoiled population had a mean crossing average of 2.5 ± 1.4 compared to the nicked population which was significantly lower at 0.9 ± 1.0 (p=2.8×10^−9^, ****) (**Figure 6F)**.

**Figure 6:**
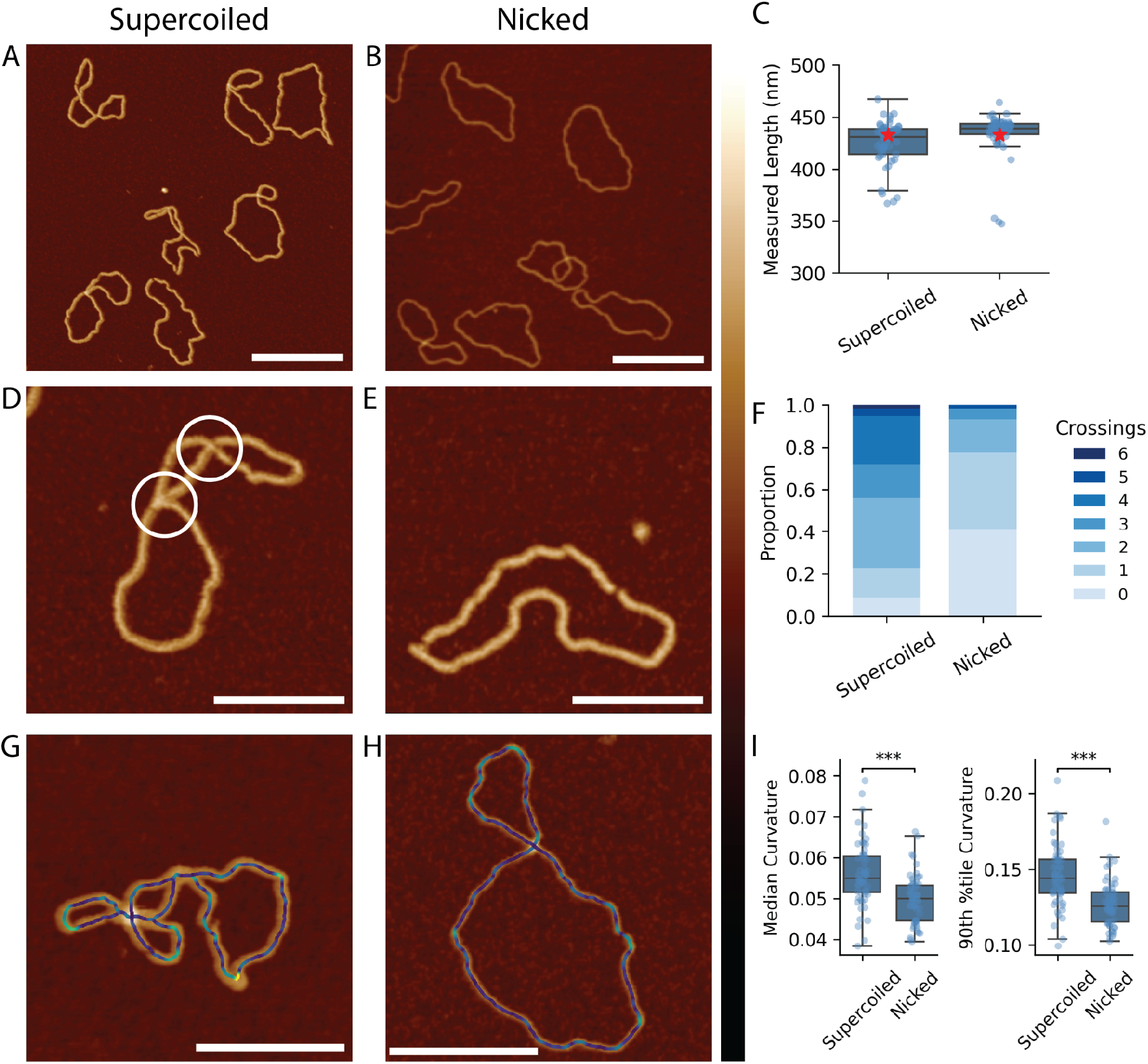
Calculation of crossing numbers allows for quantification of more subtle conformational changes. Representative zoom-out AFM images of multiple supercoiled **(A)** and nicked **(B)** pICOz DNA plasmids. Contour length box plots **(C)** for supercoiled and nicked datasets (expected pICOz length: 433 nm, indicated by red star). Representative zoom-ins of a supercoiled **(D)** and a nicked **(E)** pICOz plasmid. Stacked bar charts displaying the proportion of numbers of crossings in a plasmid between supercoiled and nicked pICOz plasmid datasets **(F)**. Scale bars: A, B 160 nm, D, E 80 nm. Height scale −3 nm to +4 nm. Statistical significance for the median curvature was determined using a Student’s t-test and for 90th percentile of curvature, a Mann-Whitney, non parametric t-test. ***: p ≤ 0.001.

#### 2.7.2. Curvature analysis shows differences in molecular conformation between supercoiled and nicked plasmids

Understanding how DNA flexibility responds to changes in supercoiling requires metrics that capture local curvature variation along the molecular backbone. Here the curvature is calculated by a rate of change of direction of the spline, with quick changes in direction measured as high curvature, and slow changes in direction as low curvature (**Figure 6G,H**). Curvature statistics calculated for supercoiled and nicked plasmids showed that nicked plasmids exhibit a significantly reduced median curvature (0.056 ± 0.008) and 90th percentile of curvature (0.15 ± 0.02) compared to the supercoiled plasmids (median = 0.050 ± 0.006, p=4.3×10^−6^,***. 90th percentile = 0.13 ± 0.02, p=2.1×10^−7^,***) (**Figure 6I**). The reduced median curvature indicates that nicked plasmids adopt a more uniform and smoothly curved conformation and the lower 90th percentile demonstrates a lack of local curvature maxima, consistent with the more open and circular structures observed in the AFM images. Conversely, the curvature statistics of the supercoiled plasmids describe conformations with tighter global and local bends, reflecting their torsionally constrained nature. These results demonstrate the utility of this method for generating accurate quantitative statistics and reliably distinguishing between different DNA topological states at the single-molecule level.

### 2.8. Changes in sequence-driven plasmid flexibility are described by curvature metrics

These advanced tracing statistics can be further leveraged to explore how changes in DNA sequence impact molecular conformation. Using two variants of the pICOz plasmid, pICOz with 12 repeats of the telomeric insert, TTAGGG (pTEL12, **Figure 7A**) and pICOz with an AT rich insert (p3AT, **Figure 7B**), we were able to quantify how changing the local GC composition of the plasmids affected its flexibility and shape (n=67 telomeric insert, n=41 AT rich insert). The 72-base pair telomeric insert in the pTEL12 has a GC content of 50%, whereas the AT rich insert of the p3AT plasmid measures 100 base-pairs and comprises only 14% GC. GC pairs form 3 hydrogen bonds between opposing bases compared to the 2 bonds formed between AT pairs, which contributes to increased structural rigidity of the DNA helix. The increased rigidity of the telomeric sequence due to the high GC content is reported to reduce the ability of the helix to compensate for torsional strain through change in twist^42^. Consequently, when negative supercoiling is introduced into the helix, the GC-rich telomeric region may resist this underwinding. Unable to accommodate increased torsional strain as twist, to minimise the energetic cost within the constrained pTEL12 plasmid, it is plausible that a greater proportion of the torsional strain is partitioned into writhe resulting in an increased number of crossings 3.3 ± 1.7 **(Figure 7D,F**) and corresponding higher median (0.06 ± 0.01) and 90th percentile curvature (0.15 ± 0.02) **(Figure 7G,I**). In contrast, the p3AT plasmid, which contains a greater proportion of AT base pairs in the same region, may be more flexible and therefore able to effectively accommodate changes in torsional strain through change in twist **(Figure 7E,F**). The p3AT plasmids exhibit significantly fewer self-crossings (1.9 ± 1.5, p=9×10^−6^,***) and a significantly smoother curvature profile (median=0.05 ± 0.01, p=0.0003,***, 90th percentile=0.15 ± 0.02, p=0.02,*) compared to the pTEL12 plasmid.

**Figure 7:**
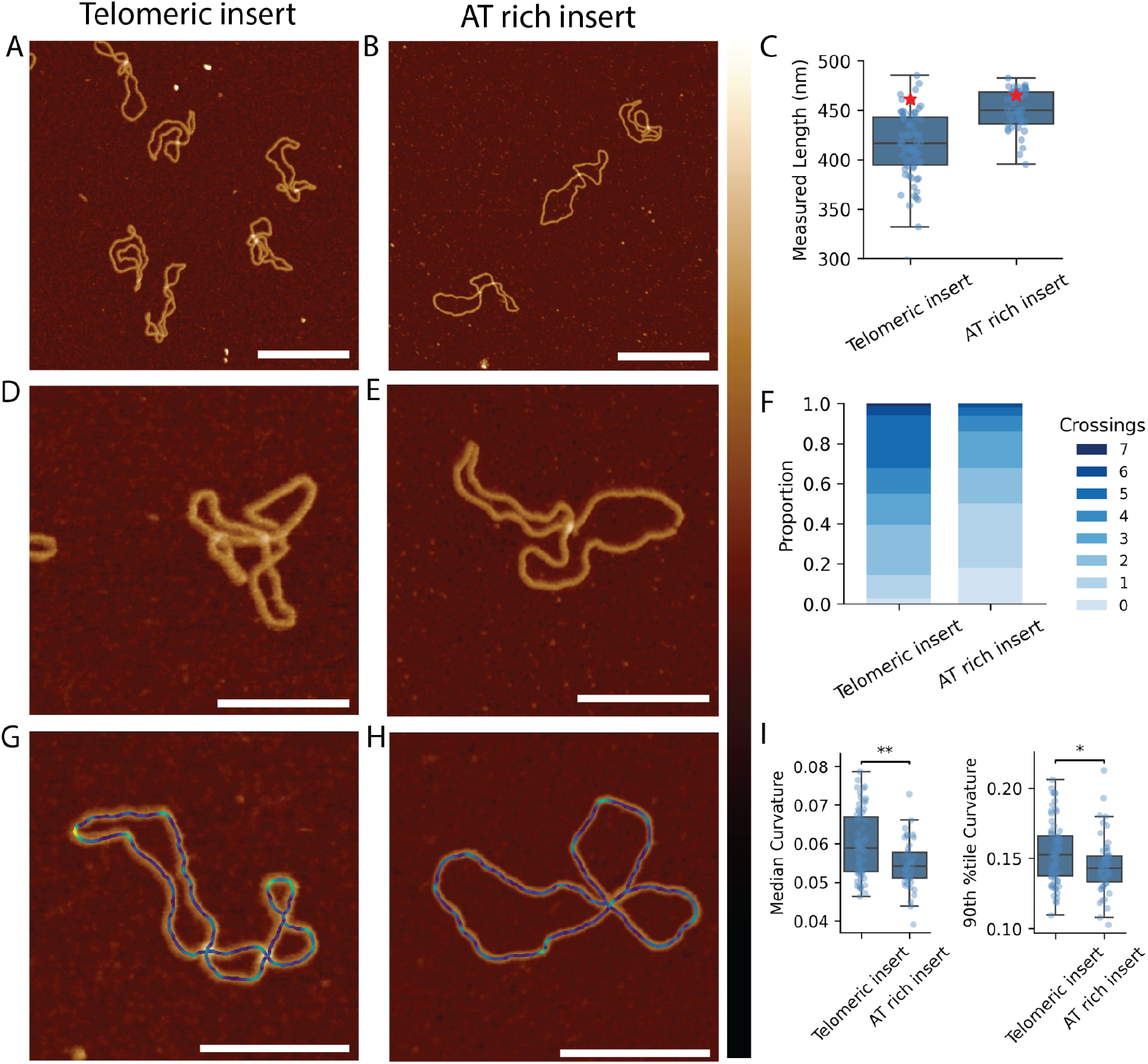
Using TopoStats to measure the effect of DNA sequence on plasmid conformation. AFM images of DNA plasmids with telomeric, **(A)** and AT-rich, **(B)** inserts. A comparison of the measured lengths of the two plasmid types, **(C)** with the red stars indicating the expected length of each plasmid. Representative zoom-ins of the plasmids **(D), (E)**. Comparison of the number of self-crossings present in the two plasmid types, **(F)**. Visualisations of curvature intensity for representative plasmids of both types **(G), (H)**. A comparison of the median and 90th percentile of curvature values for both sample types, **(I)**. Scale bars: zoom outs = 160 nm, standard crops = 80 nm, curvature crops = 80 nm. Height scale: −3 to 4 nm. Statistical significance for the median curvature was determined using a Mann-Whitney, non-parametric t-test, and for 90^th^ percentile of curvature, a Student’s t-test. **: p ≤ 0.01. *: p ≤ 0.05.

Given that both plasmids are of a similar length (pTel=1356 bp and p3AT=1367 bp, **Figure 7C**) they should have the same level of native supercoiling (ΔLk = −8) partitioned between twist and writhe. In the absence of sequence dependent effects, we would therefore expect both the pTEL12 and p3AT plasmids to have the same measured contour length, global topology and thus adopt similar surface conformations e.g., the same number of crossings and curvature. However, despite comprising only a small fraction of the total plasmid length (5% for pTel12, 7% for p3AT), the change in number of crossings and relative curvature suggest subtle sequence differences may influence the twist-writhe partitioning, and therefore global conformation of the plasmid. Taken together these data are consistent with the hypothesis that AFM, combined with advanced and accurate molecular measurements, can detect and quantify sequence dependent changes in plasmid conformation of a surface. More broadly, this highlights the power of TopoStats for studying structural effects of DNA-binding proteins, enzymatic activity and other complex systems influencing DNA topology.

## 3. Discussion

This upgraded version of TopoStats retains the core functionality of the original TopoStats software including image loading and preprocessing, object segmentation, statistical extraction and data export, but has been expanded to introduce several new functionalities. Through the development and integration of AFMReader we have removed instrument-specific barriers, enabling loading and analysis of AFM images from multiple platforms with varied file formats. We have enhanced image preprocessing to include scar removal and custom row alignment to ensure proper flattening, which is crucial for obtaining accurate statistics downstream. To further improve object segmentation, we have implemented deep-learning segmentation into the TopoStats workflow. By making it possible for user-defined TensorFlow deep-learning models to be imported as part of the segmentation pipeline, we can handle even more complex and heterogeneous biomolecular systems.

Our new tracing pipeline refines conventional skeletonisation algorithms by leveraging the unique height information in AFM data to accurately align skeletons to molecular backbones. This allows the generation of accurate height-traces to extract information such as the helical periodicity of a molecule. TopoStats overcomes the typical difficulties with tracing through strand intersections by pairing emanating branches, enabling over- and under-passing strands to be distinguished. We demonstrated the efficacy of the tracing pipeline using DNA as a model system showing that the workflow is able to accurately trace single DNA molecules and is capable of resolving overlapping molecules resulting from random surface adsorption. This functionality enables the technique to serve as a data-cleaning step, lessening the reliance on meticulous sample preparation to prevent molecular overlap and enabling a greater number of relevant molecules to be retained for analysis. The potential of the tracing pipeline is further enhanced by the addition of crossing and curvature analysis which provides insight into global and local changes in molecular shape. We have shown that nicked and supercoiled plasmids can be distinguished by quantifying changes in curvature and writhe, and these metrics can be used to probe sequence dependent changes in plasmid conformation and flexibility. Beyond DNA, these tools will enable users to explore diverse samples including polymers and filaments, proteins, globular structures and membranes, extracting rich structural data.

TopoStats represents a significant step towards making quantitative analysis more accessible to the AFM community, advancing a field that remains comparatively underdeveloped relative to other imaging techniques such as optical microscopy. In developing TopoStats, we placed strong emphasis on software design, usability, and reproducibility, guided by the FAIR4RS principles to ensure the software is findable, accessible, interoperable, and reusable. Findability is supported through public hosting of the software on GitHub, PyPi, and an online documentation site, making each version easily discoverable, citable, and accessible to the community. Accessibility is enhanced through open-source availability and comprehensive documentation covering every stage of the TopoStats workflow, which is automatically generated, versioned, and published online whenever new functionality is added to ensure up-to-date guidance for both users and developers. Interoperability is achieved through the use of open, standardised data formats (CSV, HDF5, PNG, SVG, TIFF), enabling results to be easily visualised and integrated with other analytical tools. Reusability is supported by robust software-engineering practices, including automated testing throughout the codebase to ensure reliability and consistent functionality during development, a modular architecture that supports batch processing and multi-core parallelisation for efficient large-scale analysis, and a unified YAML configuration file that contains all processing parameters. Metadata describing the software version, configuration parameters, and analysis conditions are stored alongside results, ensuring transparency and full reproducibility. Designed with user needs in mind, TopoStats fosters community engagement, allowing users to suggest new file-format support, contribute novel analytical modules, and help shape future developments through its open GitHub repository. In doing so, TopoStats lays the groundwork for a more open and collaborative, quantitative AFM research.

Collectively, these developments establish TopoStats as an advance for quantitative AFM image analysis, offering a unified, open, and reproducible platform that greatly expands the analytical scope and accessibility of this technique. The extensive toolbox included in TopoStats enables users to extract detailed structural information from their images bridging the gap between AFM imaging and quantitative molecular analysis.

## 4. Methods

All code for TopoStats, including its development history is publicly available on GitHub (https://github.com/AFM-SPM/TopoStats). Documentation is hosted via its documentation website (https://afm-spm.github.io/TopoStats/main/index.html). The main methods and algorithms implemented in TopoStats are detailed in the Methods section, and all configurable parameters are listed and described in **Supplementary Table 2**.

### 4.1. Image loading

AFM images are loaded using our AFMReader software (GitHub: https://github.com/AFM-SPM/AFMReader; Documentation website: https://afm-spm.github.io/AFMReader/). AFMReader incorporates a mixture of community-built packages and in-house decoding scripts for the loading of its various supported file formats (**Supplementary Figure 1**). Images in binary formats are loaded using Python and decoded according to their formats. AFMReader loads images and extracts the image data, and the pixel size in nanometres, which allows for the conversion between pixels to real world length. Occasionally with some formats, the pixel size is not stated but instead must be calculated from available metadata for the size and resolution of the image. Once these quantities have been obtained, they are stored in a unified format and returned to the calling script, able to be used interchangeably regardless of file type.

### 4.2. Image preprocessing

Raw AFM images undergo a series of preprocessing stages to remove imaging artefacts ahead of downstream analyses. Firstly, scan-line artefacts are corrected by subtracting the median height of each scan row to eliminate row-wise z-offsets and align the background to a uniform zero height. Tilt removal is achieved by fitting and subtracting a plane based on the median heights of each row and column. A second-order polynomial is then fitted to the row medians and subtracted to correct parabolic curvature. Finally, a fit of the form *z* = *a* * *x* \**y* is applied to remove saddle-shaped distortions. After these corrections, the overall image median is subtracted to restore the background to zero height. When many foreground objects are present, flattening becomes less accurate because median-based background fitting fails. To address this, the image is re-flattened while masking out foreground regions, identified through height-based thresholding. Height masks are binary arrays distinguishing background and foreground pixels. Three thresholding options are available: Otsu, standard deviation, and absolute. Otsu thresholding automatically determines a threshold for bimodal height data, scaled by a user-defined factor. Standard deviation thresholding sets the threshold as the mean height plus a user-defined multiple of the standard deviation. Absolute thresholding uses a fixed, user-specified height value in nanometres. Finally, to reduce high-gain noise in the image, a Gaussian filter is applied with user-configurable strength, set to 1 pixel by default.

### 4.3 Object segmentation

#### 4.3.1 Traditional segmentation

Objects of interest are identified using height-based masking, as in the flattening stage. The image is thresholded according to one of three user-configurable methods, Otsu, standard deviation, or absolute, each independently adjustable from the flattening thresholds. The resulting binary object mask can optionally exclude any objects touching the image edges, as these cannot be measured reliably due to incomplete visibility. Objects that are either too large or too small can also be removed based on user-defined area thresholds (in nm^2^), with mask pixels outside these limits set to zero.

#### 4.3.2 Deep learning segmentation

After initial object detection, a deep-learning model can optionally be applied to refine segmentation and improve accuracy. Each detected object is cropped from the image, and a user-defined padding is added around the bounding box to adjust the relative scale of the object. The crop is then converted into a square by expanding the shorter dimension of the bounding box, while ensuring it remains fully within the image boundaries. This process produces square image patches centred on each object of interest, suitable for deep-learning-based re-segmentation. The deep learning model is loaded from a user-specified path, and each identified object is re-segmented using a normalised image crop. For each crop, pixel height values are first clipped to user-defined lower and upper normalisation boundaries. The lower boundary is then subtracted, and the result divided by the difference between the upper and lower boundaries, rescaling all values to the range 0-1. This normalisation ensures input consistency required for deep learning inference. Finally, each crop is resized to match the input resolution expected by the loaded model. The model then generates a refined segmentation for each object. The batch and channel dimensions are removed, and the floating-point prediction is converted to a binary mask using a user-defined sensitivity threshold (default 0.5), corresponding to the probability at which the model is more confident that a pixel belongs to the foreground than the background. The resulting segmentation mask is then resized to match the original crop dimensions. Before reintegration into the overall object mask, the segmentation result is optionally checked for any newly introduced regions that are disconnected from the original threshold-based segmentation. This step prevents erroneous inclusion of nearby, unrelated objects. A new blank binary mask is then populated with the validated segmentations and used in all subsequent analysis steps.

### 4.4. Close end joining

Objects of interest often contain low regions of single-stranded DNA that can cause the binary masks to be split up into multiple, disconnected masks. We implemented a simple algorithm to try to identify and connect the mask ends, crossing the gaps to produce a full mask again. This process starts with producing a skeleton of the object mask. This skeletonisation process is identical to the main skeletonisation process used later on in the pipeline, where the mask is smoothed, skeletonised using height-based weighting and pruned of spurious branches. After a skeleton has been produced, “endpoints” and “junctionpoints” are identified via convolution. “Endpoints” are points that are at the ends of skeleton branches and have only one pixel neighbour. “Junctionpoints” are points that reside in the intersections of multiple branches and have more than two neighbours. Each point is compared to all other points, the distances between them are calculated and if within a user-configurable distance, flagged as potential connections. Many points may be within range of each other, so groups of candidates are created, and then once all groups have been constructed, they are individually considered. If there are two endpoints and zero “junctionpoints” in the group, then these are simply connected to each other. If there are four endpoints and zero “junctionpoints” in the group, then an algorithm searches for any hard-connected “endpoints” - ones that are already connected by the skeleton. The aim of this is to identify two “endpoints”, that are not directly connected via skeleton to any others in the group, as these are likely to be the entry and exit points for the group. An algorithm follows the skeleton from each “endpoint” and stores the “endpoints” that are connected to each other. If there are exactly two “endpoints” that are not connected, then processing can continue, else, the case is discarded. Assuming two “endpoints” are not connected via skeleton, labelled points A and D, two pathfinding options are produced, using Dijkstra’s algorithm with a cost based on the inverse of the height map to produce best routes: ABCD, and ACBD. The shortest route is then taken, as it is assumed that it’s less likely for the separated fragment to be significantly rotated. If there is exactly one “endpoint” and one “junctionpoint” in the group, then the “endpoint” and “junctionpoint” are connected. Dijkstra’s algorithm, using a cost map of the inverse height of the image is used to produce a path between points to be connected, then, a measure of the average width of the mask is made, and the single-pixel path is dilated until it matches the average width of the mask, connecting the points together.

### 4.5. Obtaining statistical descriptors for objects of interest

The spatial morphology of each segmented object is quantified using a suite of geometric and height-based metrics (**Supplementary Figure 3**). When applicable, measurements in pixel units are converted to nanometres by multiplying by the pixel-to-nanometre scaling factor the appropriate number of times (once for length, twice for area, thrice for volume). Each object is assigned a unique identifier (grain_number), corresponding to the order of indexing within the full image segmentation mask.

The object’s centre coordinates are computed as the mean of all pixel coordinates within its mask. Radius statistics are derived from the set of Euclidean distances between the centre and each edge pixel, while height statistics are calculated from all pixel height values in the mask. The object area is defined as the number of pixels within the mask, and the volume as the sum of their height values. The Cartesian bounding box area corresponds to the area of the smallest axis-aligned bounding box containing the object. In contrast, the smallest bounding length, width, and area refer to the minimum dimensions of the tightest bounding box independent of image orientation. To compute this, a convex hull is generated using the Graham Scan algorithm, and bounding boxes are iteratively constructed by rotating each hull simplex to the x-axis via a rotation matrix. From these, the smallest bounding area and dimensions are identified. Minimum and maximum Feret diameters are obtained similarly by iterating over each hull simplex and measuring the distances between opposing hull edges. All calculated statistics are stored and exported in CSV file format for further analyses (**Supplementary Tables 3 and 4)**.

### 4.6. Strand tracing

Strand tracing consists of three main stages: skeletonisation for identifying branches, crossing analysis for determining the route strands take through crossings, and vectorisation which reconstructs complete molecular traces from this information.

#### 4.6.1 Skeletonisation

Before skeletonisation, the object mask is smoothed to reduce jagged edges that can produce spurious branches not representative of the true structure. Two smoothing methods are available: dilation and Gaussian. Dilation performs a user-configurable number of morphological dilations, thickening and smoothing the mask edges, while Gaussian smoothing applies a Gaussian filter followed by thresholding at 0.5 to restore a binary mask. If both are enabled, they are applied in parallel, and the method that alters the fewest pixels relative to the original mask is retained. To preserve internal structure, holes are tracked during smoothing; any hole removed can optionally be reinserted if its area lies within user-defined thresholds. Skeletonisation is performed using a modified Zhang algorithm^38^ that incorporates height information. In each iteration, the pixels marked for removal are ordered by their height values, and only the proportion of pixels below a user-specified height percentile are removed. For example, a threshold of 0.4 removes only the lowest 40% of pixels by height, biasing the skeleton toward higher regions of the image. Branches, crossings, and endpoints are identified by convolving the binary skeleton with a 3×3 kernel of ones.

This produces a map in which each skeleton pixel is assigned a value corresponding to the number of connected neighbours. Pixels with two neighbours are classified as branches, those with three or more as junctions, and those with one as endpoints. Spurious branches are then removed in a pruning step. Junction pixels (those with ≥3 neighbours) are temporarily removed, isolating the branches, which are then analysed individually. Branches may be pruned if they are shorter than a user-defined minimum length or if their height statistics (minimum, median or midpoint height) fall below a configurable threshold. There is also an option to set whether to only prune branches by height if they contain an endpoint.

Skeleton statistics are then computed using the Skan library^43^, which indexes each branch, its connected branches, and its classification (e.g. endpoint-to-endpoint, junction-to-endpoint, junction-to-junction, or loop). Branch lengths, measured in pixels by Skan, are converted to nanometres and summed to obtain total branch length. The number of endpoints and junctions are also recorded. The average strand width is calculated by performing a distance transform on the object mask, sampling the resulting distances along the skeleton, and averaging them. All measurements are stored as part of the object’s statistics and exported to CSV.

#### 4.6.2 Crossing analysis

Crossing analysis identifies and characterises regions where strands overlap within the skeletonised structure. The process involves four main steps: detection of crossing regions, correction of fragmented crossings, branch pairing, and determination of strand stacking order. Crossing regions are first identified by locating junctions, points where three or more branches of the skeleton meet, and merging those that lie within a user-defined distance. For DNA data, a distance of approximately 6-8 nm is typically appropriate, corresponding to roughly twice the apparent DNA width of 2 nm. The algorithm iterates over each branch extending from a junction and traverses it for the specified distance; if another junction is encountered within this range, the connecting pixels are flagged to mark them as part of the same crossing region. In some cases, a true crossing involving an even number of strands is erroneously split into multiple junctions containing an odd number of branches. To correct this, crossing regions that exhibit odd branch counts and lie within a user-specified distance are merged. The number of emanating branches for each junction is determined by counting adjacent pixels connected to the skeleton.

Each crossing region is analysed to determine how branches enter and exit. For every branch connected to a crossing, a user-defined segment length is traced and averaged to form a representative vector from the crossing centre toward the branch. Branches are then paired based on the alignment of their vectors. Two pairing strategies are available: (1) iterative pairing of the most closely aligned branches until none remain, or (2) exhaustive pairing evaluation, selecting the combination with the lowest total alignment error. In cases with an odd number of branches, the user can specify whether pairing should proceed or be skipped. To reduce the impact of variations between individual skeletons on the tracing output, a trace averaging step is applied. Two morphological dilations are performed on the connected branches, and the result of the first is subtracted from the second, generating two parallel traces approximately one pixel apart. These, along with the original trace, are averaged to yield a smoother and more representative strand path. The stacking order of each branch pair, indicating which strand passes over the other, is then determined. Each branch is traversed over a configurable distance, and its height profile is extracted. From the middle 40% of this segment, the maximum height is identified, and the full width at half maximum (FWHM) is computed. Restricting analysis to the central portion of the branch reduces interference from nearby crossings. Branches with larger FWHMs are interpreted as lying above those with smaller values. A confidence value is calculated as the ratio between the minimum and maximum FWHMs within each pair. At this stage, statistics including the number of crossings, mean and minimum confidence of crossings, and stacking order, are recorded and stored for subsequent analysis.

#### 4.6.3 Vectorisation

The skeleton is re-analysed using connected component analysis after temporarily removing pixels belonging to crossing regions. Each branch is linked to its corresponding stacking-order information from the crossing analysis, assigning a pseudo-height that reflects its relative position within the molecule. Tracing begins at an endpoint if present, or with the first branch index otherwise, but never within a crossing segment. The branch is followed and its coordinates added to the overall trace. When a segment ends, the adjacent segment with the highest stacking order is followed next, allowing the trace to progressively build. Each completed segment is removed from the mask to prevent retracing. The process continues until no further segments remain, with disconnected regions treated as separate molecules, resulting in vectorised traces for all objects. A simplified pseudo-trace is also generated, assigning integer pseudo-heights (0, 1, 2, …) from the stacking order and subsampling the points to fewer than 100 to reduce its complexity. This simplified trace is passed to Topoly [26] to determine the object’s topological classification and Jones polynomial, which are recorded in the object statistics. Finally, the molecule’s writhe is derived from the vectorised trace by computing cross products between adjacent branch vectors aligned along the trace direction. Overpassing and underpassing crossings are denoted by “+” and “–”, with multiple crossings within a region enclosed in parentheses. The resulting writhe string is stored in a CSV file of detailed molecular statistics (**Supplementary Table 5**).

### 4.7. Trace smoothing

Trace smoothing is applied to the vectorised skeleton traces using either spline fitting or rolling average smoothing. In the spline method, multiple B-splines are generated and averaged to produce a smoother and more representative path. The number of splines is determined by a user-defined interval, which specifies how frequently coordinates from the trace are sampled to construct each spline. For example, if the interval is set to 3 nm and trace coordinates are spaced 1 nm apart, three splines are generated, each offset by one coordinate. Splines are computed using the splprep function ^44^om SciPy^44^, with configurable parameters for smoothing strength (separately for linear and circular traces) and spline degree. The sampled splines are averaged to yield the final smoothed trace. Alternatively, rolling average smoothing applies a moving window of user-defined width to locally average neighbouring points along the vectorised trace. Following smoothing, the trace is resampled to enforce uniform point spacing. This is achieved by iterating through the coordinates, calculating distances between successive points, and interpolating or skipping points as needed to maintain consistent spacing. The resulting trace provides a smooth and evenly sampled representation of the original vectorised skeleton, suitable for downstream analysis.

### 4.8. topostats file format

The introduction of the .topostats file format (a custom HDF5-based structure) provides modularity and flexibility for users who wish to access or repurpose outputs from different stages of the TopoStats pipeline. Users can extract intermediate data such as flattened images, segmentation masks, or molecular traces for further processing, visualisation, or integration with external analytical tools. These outputs can be used directly within Python or saved as standard image files or NumPy arrays for import into common image-analysis software such as ImageJ. Our TopoStats GitHub repository provides notebooks that illustrate how to interact with .topostats files, guiding users through the extraction of relevant data and demonstrating how custom functions can be developed for bespoke analysis workflows, thereby giving users more autonomy in how they use the software.

### 4.9. AFM Sample Preparation and Imaging

8 ng of supercoiled, nicked, AT insert or telomeric insert DNA was immobilized on a freshly cleaved mica disk in 30 μL of immobilization buffer (3 mM NiCl_2_, 20 mM HEPES, pH7.4) for 5 mins. The mica was then washed four times with the same buffer, and a further 30 μL was added for imaging. All AFM measurements were performed in liquid following a previously publishe^45^protocol^45^. All experiments were carried out in PeakForce Tapping imaging mode on a FastScan Dimension XR AFM system (Bruker), using FastScan D (Bruker) probes. The PeakForce amplitude was set to 6 nm, the PeakForce Tapping frequency to 8 kHz and the PeakForce setpoints in the range: 7–15 mV, corresponding to peak forces of <70 pN. All images were taken at 0.5 nm/px at line rates of ~3–5 Hz.

The acquired images were then processed using TopoStats. Processing with TopoStats resulted in a total of n=54 supercoiled, n=55 nicked, n=62 telomeric insert, n= 47 AT rich insert molecules being retained for this analysis.

### 4.10. 3AT and Telomeric Repeat insert sequences

#### 3AT insert sequence

5’

atcctatatatatatatatatatatatatatgcgaactcatatatatatatatatatatatatatggtgccgtcatatatatatatatatatatatatat 3’

3’

taggatatatatatatatatatatatatatacgcttgagtatatatatatatatatatatatataccacggcagtatatatatatatatatatatatata 5’

#### Telomeric DNA insert

5’ ccctaaccctaaccctaaccctaaccctaaccctaaccctaaccctaaccctaaccctaaccctaaccctaa 3’

3’ gggattgggattgggattgggattgggattgggattgggattgggattgggattgggattgggattgggatt 5’

## Data availability statement

All data used in this publication, and plotting scripts [will be] publicly available on Figshare upon publication. Plasmids used in the study will be deposited on AddGene upon publication.

## Code availability statement

All code developed in this publication is publicly available on the main GitHub branch of TopoStats https://github.com/AFM-SPM/TopoStats.

## Contributions

Formal contributions in authorship order (CrediT taxonomy):

Conceptualization: A.L.B.P;

Data curation: S.W, T.A.F, M.G, L.W, T.A, N.S;

Formal Analysis: S.W, T.A.F;

Funding acquisition: A.L.B.P;

Investigation: A.L.B.P, L.W, S.W, M.G, T.A.F, T.E.C;

Methodology: S.W, M.G, L.W, N.S, A.L.B.P;

Project administration: A.L.B.P, T.E.C;

Resources: A.L.B.P;

Software: S.W, M.G, N.S, L.W, T.A, A.L.B.P;

Supervision: A.L.B.P, L.W, T.E.C;

Visualization: S.W, T.A.F, T.E.C;

Writing – original draft: S.W, T.A.F;

Writing – review & editing S.W, T.E.C, T.A.F, L.W, M.G, A.L.B.P:

## Acknowledgements

The authors would like to thank Professor Jens Staal for designing the original pICOz cloning vector used in these analyses, and Dr Matthew D. Newton and Dr Ellen G. Allwood for generating and kindly providing the pICOz_Tel12 variant. A.L.B.P. acknowledges funding from UKRI Future Leaders Fellowship MR/W00738X/1, the Henry Royce Institute for Advanced Materials, funded through EPSRC grants EP/R00661X/1, EP/S019367/1, EP/P02470X/1, and EP/P025285/1 and Xinyue Chen for Dimension FastScan access and support at Royce@Sheffield. S.W. was supported by the Medical Research Council (MR/W006944/1)

## Notes

### Competing Interest Statement

The authors have declared no competing interest.

