## Supplementary Information for "Deep Learning-Enhanced TopoStats for the Automated Quantification of DNA and Complex Biomolecular Structures"

| File format name | Description |
|---|---|
| **Supported file formats** | |
| .spm | Bruker primary format |
| .jpk | Bruker acquired format |
| .jpk-qi-image | Bruker QI imaging mode |
| .h5-jpk | Bruker open acquired format |
| .gwy | Gwyddion file format |
| .ibw | WaveMetric format |
| .stp | WSXM AFM format |
| .top | .stp format variant |
| .topostats | TopoStats HDF5-encoded output |
| .asd (partial) | High speed AFM format |
| **Unsupported file formats** | |
| .nhf | Nanosurf format |
| .aris | Imaris Oxford Instrument format |

| | |
|---|---|
| .tiff | Common image format |

**Supplementary Table 1 .** AFM file formats currently supported in TopoStats2 via AFMReader.

| Section | Parameter | Default Value | Description |
|---|---|---|---|
| **General** | | | General options |
| | base_dir | ./ | Directory containing the data to process |
| | output_dir | ./output | Directory to output results to |
| | cores | 2 | Number of CPU cores to utilise |
| | file_ext | .spm | File type of the data files to search for and load |
| **Loading** | | | Options pertaining to loading AFM data |
| | channel | Height | Image channel to load from the data file |
| **Filter** | | | Options pertaining to flattening & preprocessing |
| | row_alignment_quantile | 0.5 | Quantile to assume the background value is at. Lower values may produce better flattening in images with more data above the surface |
| | threshold_method | std_dev | Method to use for thresholding objects from the background |
| | otsu_threshold_multiplier | 1.0 | Multiplier to apply to the calculated Otsu threshold |
| | threshold_std_dev | 1.0 | Number of standard deviations from the mean to use as the height threshold |
| | threshold_absolute | 1.0 | Absolute height threshold in nanometres |
| | gaussian_size | 1.0 | Amount of pixels of gaussian smoothing to apply to the flattened image |
| | remove_scars/run | false | Whether to interpolate over scars in the image |
| **Grains** | | | Options pertaining to identifying objects of interest |
| | grain_crop_padding | 1 | Number of pixels to pad the grain's bounding box by |
| | threshold_method | std_dev | Method to use for thresholding objects from the background |
| | otsu_threshold_multiplier | 1.0 | Multiplier to apply to the calculated Otsu threshold |

| | | | |
|---|---|---|---|
| | threshold_std_dev | 1.0 | Number of standard deviations from the mean to use as the height threshold |
| | threshold_absolute | 1.0 | Absolute height threshold in nanometres |
| | area_thresholds | 300, 3000 | Lesser and greater area thresholds in nm^2 below / above which grains are removed |
| | remove_edge_intersecting_grains | true | Whether to remove grains that intersect the image bounds |
| **unet_config** | | | Options pertaining to the re-segmentation of identified objects of interest using a user-provided, pre-trained TensorFlow deep-learning model |
| | model_path | null | System path to a trained TensorFlow model |
| | upper_norm_bound | 5.0 | Pre-segmentation upper normalisation bound |
| | lower_norm_bound | -1.0 | Pre-segmentation lower normalisation bound |
| | remove_disconnected_grains | false | Whether to remove regions in predicted masks that do not intersect the original mask |
| | confidence | 0.5 | The confidence threshold for the binarisation of the prediction |
| **disordered_tracing** | | | Options pertaining to skeletonisation |
| | mask_smoothing_params/gaussian_sigma | 2 | Amount of gaussian smoothing to use when smoothing a mask before skeletonisation |
| | mask_smoothing_params/dilation_iterations | 2 | Number of morphological dilation processes to run when smoothing a mask before skeletonisation |
| | mask_smoothing_params/holearea_min_max | [0, null] | The minimum and maximum size in nm^2 of holes in the mask pre-smoothing to re-add after smoothing |
| | skeletonisation_params/height_bias | 0.6 | Percentage of pixels to remove from the candidate pixels, ordered by height. Lower value results in stronger biasing towards high regions, higher values approach a Zhang-style skeletonisation |

| | | | |
|---|---|---|---|
| | pruning_params/max_length | 10 | Maximum length in nm to remove a branch that does not connect to another branch |
| | pruning_params/height_threshold | null | The height below which to remove branches |
| | pruning_params/method_values | mid | The method to obtain a branch's height for determining if it should be removed |
| | pruning_params/method_outlier | mean_abs | The method for determining outlier heights of branches |
| | pruning_params/only_height_prune_endpoints | false | Whether to only apply height based branch removal to branches that do not connect to other branches |
| **nodestats** | | | Options pertaining to crossing analysis |
| | node_joining_length | 7.0 | Distance in nm to connect nearby crossing points |
| | node_extend_dist | 14.0 | Distance in nm to join nearby odd-branched crossings |
| | branch_pairing_length | 20.0 | Distance in nm to follow a branch to collect height data for crossing analysis |
| | pair_odd_branches | false | Whether to try to pair branches of crossings with odd-numbers of branches |
| **splining** | | | Options pertaining to smoothing & splining |
| | method | rolling_window | Whether to use splining or rolling window averaging for trace smoothing |
| | rolling_window_size | 20.0e-9 | Size of the rolling average window, in metres |
| | rolling_window_resampling | true | Whether to resample rolling-window averaged points to ensure they are equally spaced along the trace |
| | rolling_window_resample_regular_spatial_interval | 0.5e-9 | The distance in metres that resampled traces' points should be spaced |
| | spline_step_size | 7.0e-9 | The sampling rate in metres of the spline(s) |
| | spline_linear_smoothing | 5.0 | The smoothing for splines that do not connect to themselves to form a |

| | | | |
|---|---|---|---|
| | | | loop |
| | spline_circular_smoothing | 5.0 | The smoothing for splines that do connect to themselves to form a loop |
| | spline_degree | 3 | The degree of the spline polynomial |
| **plotting** | | | Options pertaining to plotting of results |
| | savefig_format | null | What format to save images in, either PNG, SVG, PDF |
| | savefig_dpi | 100 | The DPI for each saved output image |
| | grain_crop_size_nm | -1 | The size of the crops in nm for grains when the “grains” setting for image_set is used. Null results in tight bounding boxes |
| | image_set | core | The verbosity of output images. Core just produces the minimum. Add more modules to get detailed visual output for them |
| | zrange | Null, null | The minimum and maximum values in the colour scale for output images. Null results in using the minimum and maximum for the image |

**Supplementary Table 2 - TopoStats’ basic configuration options.** The most commonly used and generally applicable configuration options of TopoStats2, provided in a .yaml file.

| Parameter | Description |
| --- | --- |
| image | Filename of the image |
| image_size_x_m | Width of the image in metres |
| image_size_y_m | Height of the image in metres |
| image_area_m2 | Area of the image in metres squared |
| image_size_x_px | Width of the image in pixels |
| image_size_y_px | Height of the image in pixels |
| image_area_px2 | Area of the image in pixels squared |
| grains_number | Number of objects of interest found |
| grains_per_m2 | Number of objects of interest per metre squared |
| rms_roughness | Root mean square roughness of the image |

**Supplementary Table 3 - Global image statistics.** Statistics produced by TopoStats2 pertaining to each whole AFM image, allowing quantification and comparison of global metrics.

| Parameter | Description |
| --- | --- |

| | |
|---|---|
| image | Filename of the image |
| basename | Parent folder (dataset) name |
| grain_number | Index of the object of interest |
| centre_x | Centre x coordinate of the object (m) |
| centre_y | Centre y coordinate of the object (m) |
| radius_min | Minimum radius of the object (m) |
| radius_max | Maximum radius of the object (m) |
| radius_mean | Mean radius of the object (m) |
| radius_median | Median radius of the object (m) |
| height_min | Minimum height of the object (m) |
| height_max | Maximum height of the object (m) |
| height_mean | Mean height of the object (m) |
| height_median | Median height of the object (m) |
| volume | Volume of the object (m3) |
| area | Area of the object (m2) |
| area_cartesian_bbox | Area of the bounding box for the object (m2) |
| smallest_bounding_width | Length of the shorter axis of the smallest possible rectangular bounding box not aligned to cartesian coordinates (m) |
| smallest_bounding_length | Length of the longer axis of the smallest possible rectangular bounding box not aligned to cartesian coordinates (m) |
| smallest_bounding_area | Area of the smallest bounding box not aligned to cartesian coordinates (m2) |
| aspect_ratio | Ratio of the axes of the smallest possible bounding box not aligned to cartesian coordinates. |
| max_feret | Maximum feret diameter (m) |
| min_feret | Minimum feret diameter (m) |
| grain_endpoints | Number of strand end-points in the object |
| grain_junctions | Number of junction pixels in the object's skeleton |
| total_branch_length | The sum of the lengths of all branches of the object's skeleton |
| grain_width_mean | The mean width of the strands in the object |
| num_mols | The number of individual molecules contained in |

| | |
|---|---|
| | the object |
| writhe_string | A string of the under/over status of all the object’s crossing regions. |
| average_end_to_end_distance | The mean distance between the object’s strand end-points |
| total_contour_length | The total length along all smoothed ordered traces for all identified molecules |

**Supplementary Table 4 - Object statistics.** Statistics produced by TopoStats2 pertaining to each individual object in each AFM image, with a mixture of general geometric properties and more specific DNA-focused measurements.

| Parameter | Description |
|---|---|
| image | Filename of the image |
| basename | Parent folder (dataset) name |
| grain_number | Index of the object of interest |
| molecule_number | Index of the molecule within the object |
| circular | Whether the molecule contains a self-connection. |
| topology | Topological classification of the object. Ie, linear, 0_1, 3_1, etc. |
| contour_length | Length of the molecule (m) |
| end_to_end_distance | Euclidean distance between linear molecules’ endpoints |

**Supplementary Table 5 - Detailed molecule statistics.** Extra statistical information about sub-grains and molecules identified by TopoStats2, where multiple molecules or objects intersect and should be handled as a single whole object, but are identifiable as different structures.

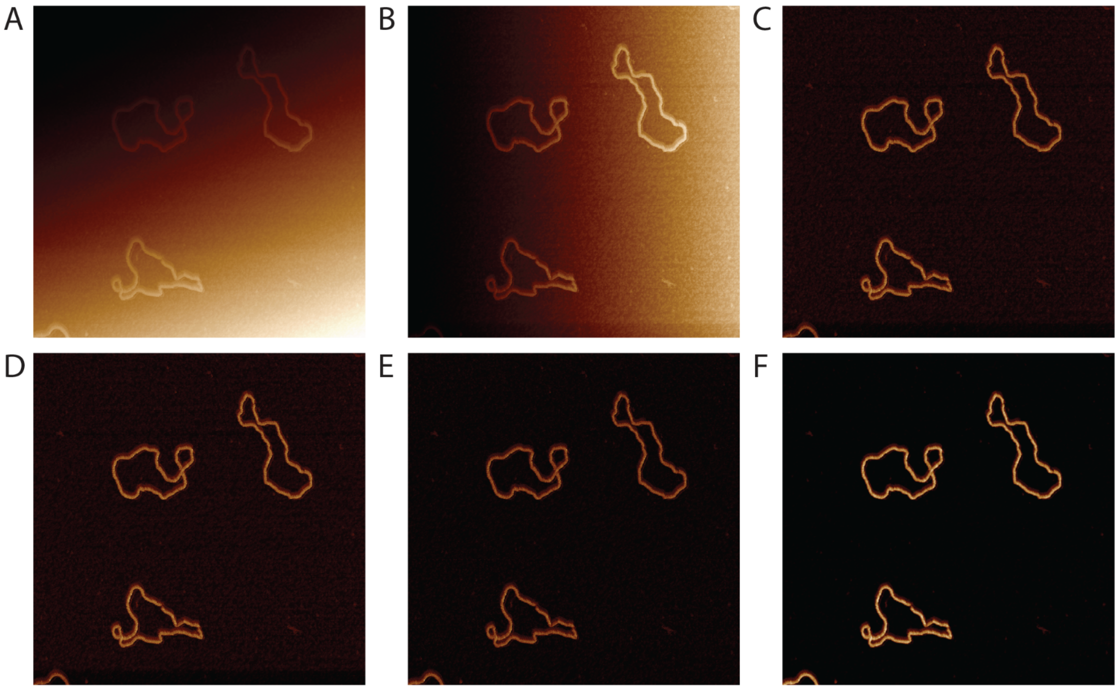


**Supplementary Figure 1. Image filtering and flattening process in TopoStats2.** Raw image **(A)**, which subsequently undergoes row alignment **(B)**, tilt removal **(C)**, and quadratic flattening **(D)**. Finally, the median height of the image background pixels are set to zero **(E),** and a user-defined Gaussian blur is applied **(F)**. Image size is 500 nm.

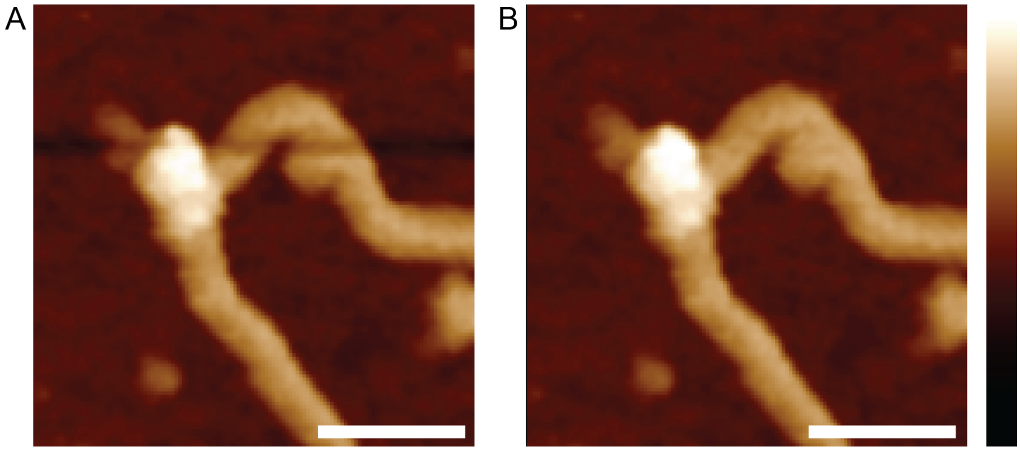


**Supplementary Figure 2. Image flattening artefacts present in TopoStats v1.0 are able to be remedied in TopoStats v2.0.** AFM image **(A)** of a DNA plasmid processed using TopoStats v1.0 method, where a flattening artefact manifests as a thick dark band across the image, erroneously lowering the height of the affected rows. The same AFM image **(B)** processed using TopoStats v2.0 method, where the flattening artefact has been remedied with use of the “row_alignment_quantile” setting being set to 0.1, resulting in no dark band, nor erroneous lowering of rows. Height scale: -3 - 4 nm. Scale bars: 20 nm.

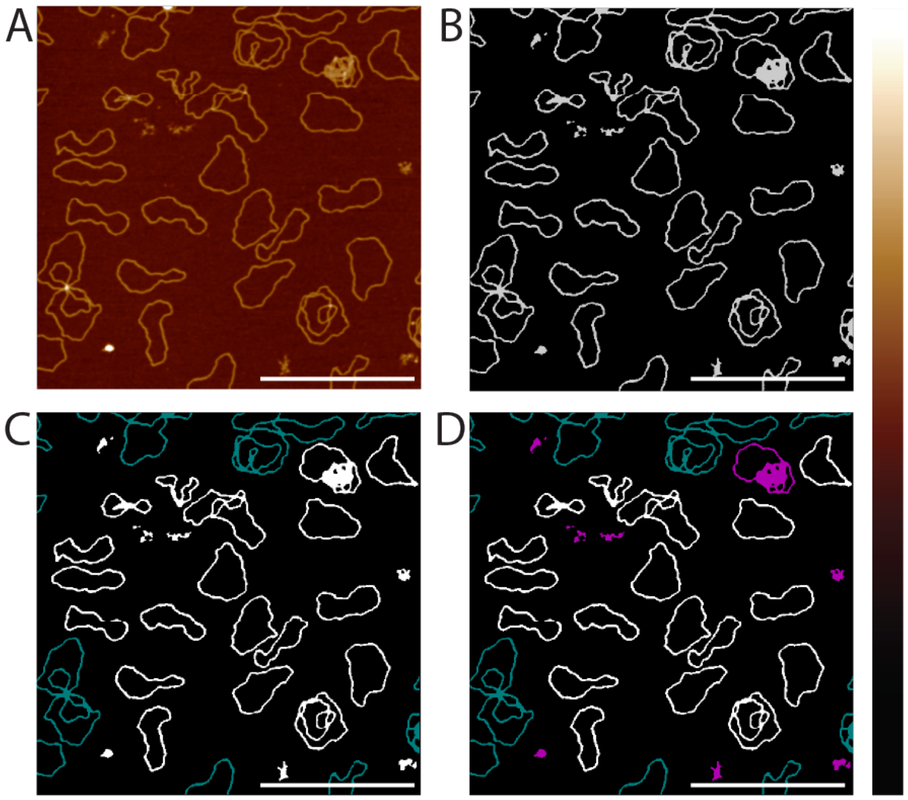


**Supplementary Figure 3. Height masking enables identification of objects of interest.** AFM image **(A)** of multiple DNA plasmids. Initial height-based binary mask **(B)** identifying objects of interest and isolating them from the background. Removal of edge-intersecting objects marked in cyan **(C)**, and objects that are too small or too large marked in magenta **(D)**. Scale bars: 400 nm.

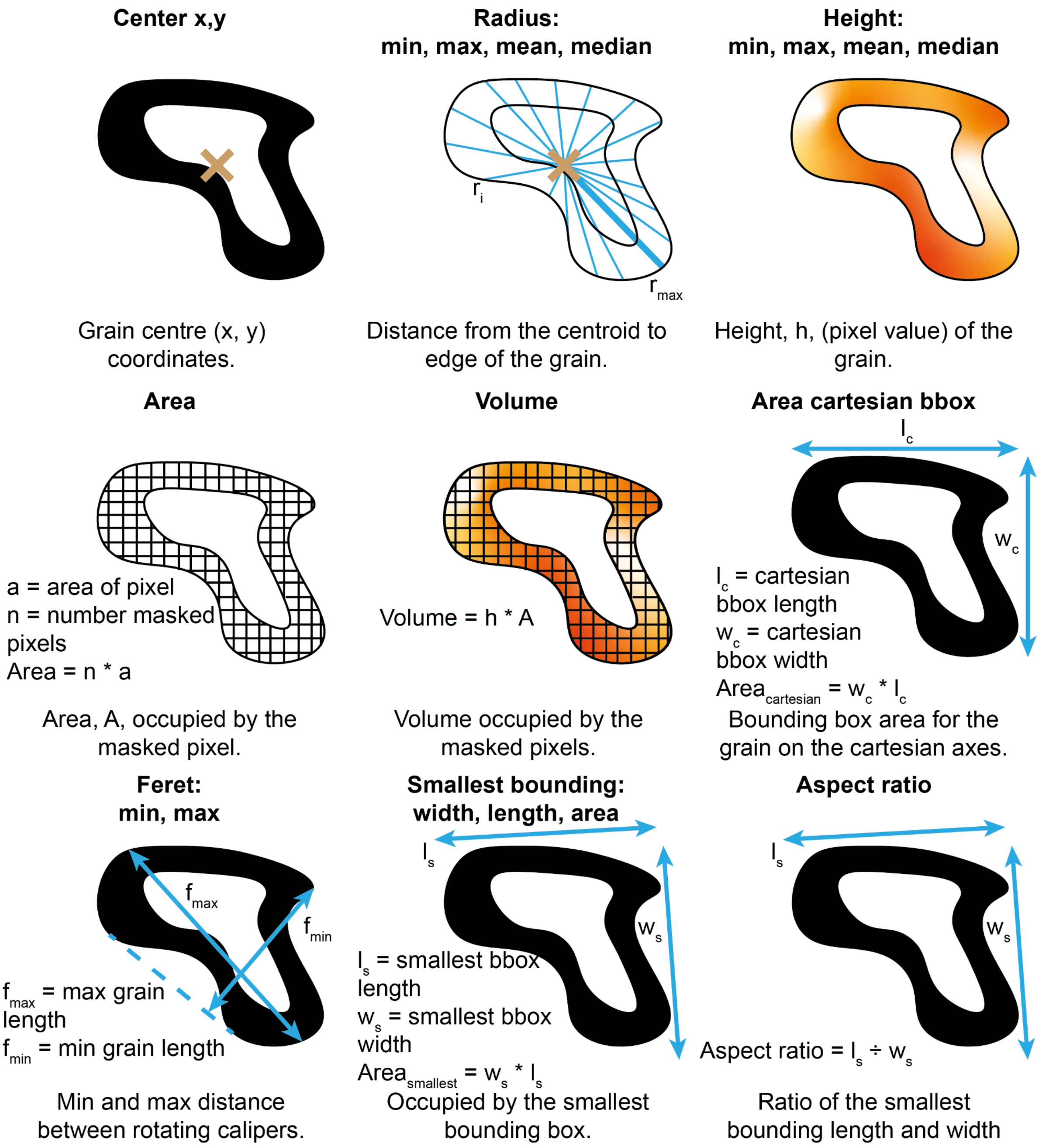


**Supplementary Figure 4: Basic grain shape can be quantified using morphological metrics.** A visualisation of the various morphological metrics that are calculated by TopoStats2 for each object of interest, and subsequently output in the results file.

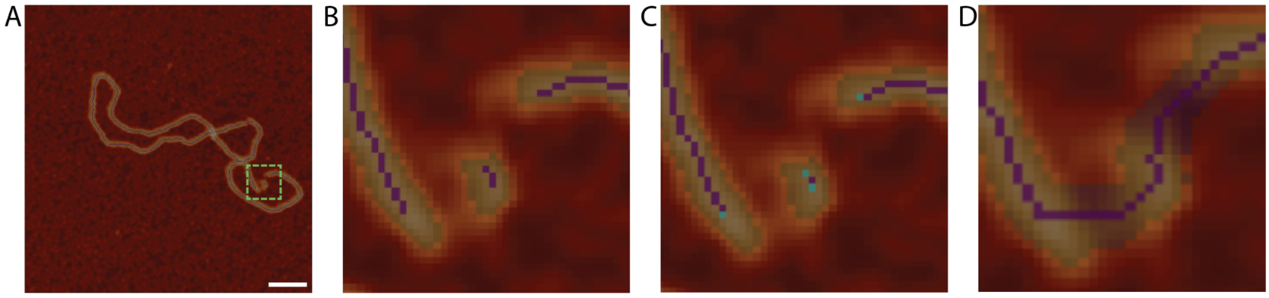


**Supplementary Figure 5: Process of end-joining to reconnect broken skeletons.**
AFM image of a pICOz plasmid with two height dips **(A)** resulting in a discontiguous mask and skeleton **(B)**. End-points are identified **(C)**, and connected via a constructed skeleton following the highest path between the endpoints and subsequent dilation to produce a filling of appropriate width, resulting in a complete mask, **(D)**.

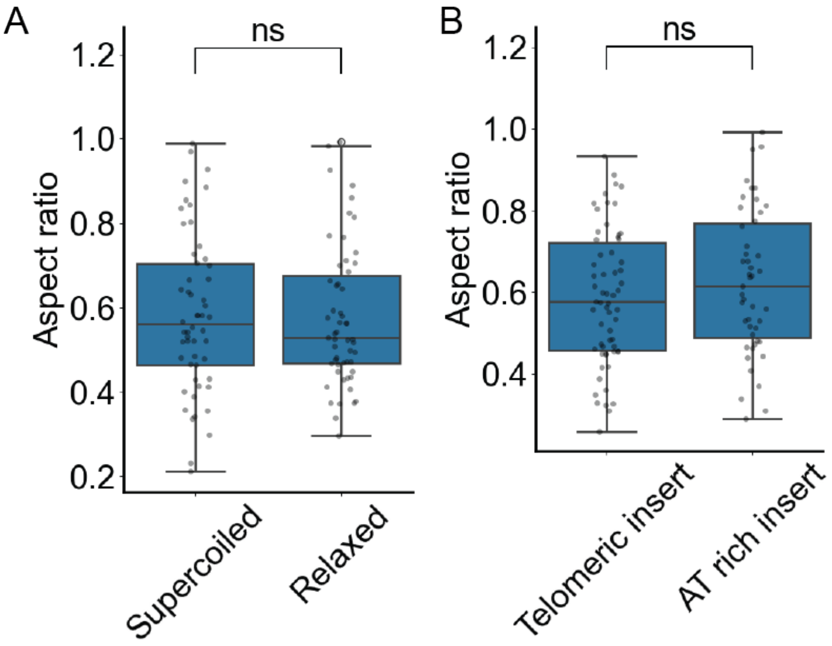


**Supplementary Figure 6: Basic morphological measurements fail to distinguish supercoiled and nicked pICOz datasets.** Comparison of supercoiled & nicked pICOz **(A)** and telomeric insert and AT rich insert **(B)** DNA plasmid datasets' aspect ratios. Statistical significance for the aspect ratio of supercoiled and nicked plasmids was determined using a Mann-Whitney, non parametric t-test, and for the telomeric insert and AT rich inserts, a Student's t-test.
